# Time Lags and Niche Shifts in a Biological Invasion of Hummingbirds

**DOI:** 10.1101/329615

**Authors:** C.J. Battey

## Abstract

Shifts in a species’ realized niche can lead to rapid population growth by increasing the carrying capacity of local habitats and allowing colonization of new areas. This process is well known in “invasive” species introduced to novel ranges by humans, but can also occur when native species expand their range. In these cases expansions may be driven either by a shift in the available environment or by a shift in the species’ use of existing niche space, but identifying the specific environmental or behavioral changes involved is often hindered by time lags in the process of colonization. Here I document a century of range shifts in the Anna’s Hummingbird (*Calypte anna*) and show that recent abundance in the Pacific Northwest is the product of a series of range and niche expansions that started in the early 20^th^ century following the spread of garden cultivation and introduced plant species in California. Demographic trends in the northwest have tracked simple models of exponential growth since populations became established in the 1960’s and 70’s, and nest records suggest that the species has delayed the beginning of the breeding season by at least 18 days in the north. Niche models trained on historic climate and occurrence data fail to predict the modern range, suggesting that climate change is not the primary cause of the expansion. Range expansions in the Anna’s Hummingbird thus closely track the dynamics of an invasive species spreading across a novel range, and were made possible by a mix of introduced plants, phenological acclimation, and an expansion of the realized climatic niche.

## Introduction

Range shifts are common on evolutionary time-scales and are widely predicted to be necessary for species’ persistence as climate change alters temperature and precipitation regimes around the world (Chen et al. 2011, Doak and Morris 2010, Keit et al. 2001, Buckley 2007). In the recent past many dramatic examples of range expansions have come from “invasive” species – typically those introduced to novel ranges by humans (Tingley et al. 2014, Veit and Lewis 1995). However, the invasion dynamics observed in introduced species can also occur when native species expand their range without direct human transport. In these cases range shifts can be caused either by an underlying change in the available environment, or by a shift in the species’ use of preexisting environmental conditions (i.e. its realized niche; Hutchinson 1957). Distinguishing these scenarios is difficult, in part because we often lack accurate data on when and where distributional changes started if human transport is not involved.

Range shifts and biological invasions are demographic processes occurring over time, so a lag between the initial colonization event and the first observation is inevitable (Kowarik 1995). Small populations may not be dense enough for population growth (Allee 1938, Veit and Lewis 1995) and are particularly vulnerable to extirpation by stochastic events (Lande 1993); creating an initial waiting period in which new regions are intermittently colonized and extirpated until one drifts high enough in abundance to begin growing stably (Kowarik 1995, Levin 1969, Hanski 1998). Once populations are established, the compounding nature of exponential growth and the imperfection of survey data mean that we often won’t observe a new species in significant numbers until its population has reached what appears to be explosive growth (Crooks 2005).

This time lag between colonization and observation means that the events that caused or allowed a range expansion – changes in climate, behavior, habitat availability, etc. – will likely occur before it is reflected in survey data. One way to incorporate this process into analyses seeking to identify the environmental or behavioral drivers of range shifts is to explicitly model population growth over time in different regions (Veit and Lewis 1995, Hanski 1998, La Sorte et al. 2007) rather than limiting analysis to a presence/absence grid. Growth rates and colonization times can then be inferred as model parameters and the timing of either initial colonization or a change in growth rates can be compared with historic climate or habitat data to ask if shifts in the environment explain distributional changes.

Here I analyze demographic and distributional data for the Anna’s Hummingbird (*Calypte anna*) in order to document the timing and population dynamics of a recent range expansion and test for evidence of a shift in the realized niche. I first use historic occurrence records to document the range of the species in the early twentieth century and map its spread to the present day. I then estimate growth rates across the range and ask if population dynamics in the native and novel ranges are qualitatively different by fitting simple demographic models to long-term survey data. To assess whether climate change can explain the distributional changes I fit niche models to historic climate and occurrence data and project these to modern climates. Last, I test for phenological acclimation in the species’ new range by comparing the timing of nest records from native and novel ranges.

### Natural History of the Anna’s Hummingbird

The Anna’s hummingbird is a member of the rapidly diversifying “Bee hummingbird” clade (McGuire et al. 2014) native to western North America. Joseph Grinnell, the early California biogeographer and systematist, described its range in 1915 as chaparral and scrub oak habitats from Baja California to the north end of the Sacramento Valley. The few observations then reported from the north coast or Klamath mountains were “doubtless beyond the regular breeding area of this species”, because “in its breeding range and throughout the year… the Anna Hummingbird adheres with remarkable closeness to the Upper Sonoran life zone” (Grinnell 1915). The species is not a classic seasonal migrant, but is known to disperse to higher elevations and latitudes after breeding (Clark and Russel, 2012).

The nesting season is unusually early, running through winter from November to May (Clark and Russel 2012, Williamson 2001). The end of the dry season (late fall) is the low point of nectar availability in most of the native range, and the first significant blooms in the chaparral – particularly manzanita (*Arctostaphylos sp.*) and currant (*Ribes sp.*) *–* start in November and peak from February to March (Jepson 1993), which may explain the early onset of nesting. Anna’s Hummingbirds are also known to nest twice in a year (Scarfe et al. 2001) with 2 eggs per nest (Stiles 1973), and an early start to the nesting season may leave more time for a second clutch.

By the early 20^th^ century, birders and naturalists had noticed Anna’s hummingbird populations increasing across the range. Grinnell and Miller (1944) and Robertson (1931) both remarked that the mid 19^th^ century introduction of Eucalyptus trees, which bloom from November to April, likely provided the food source that allowed populations to increase. Though the term was not yet in wide use by population biologists, Grinnell comes close to describing the effect as increase in carrying capacity: “This means that the rigors of a ‘minimum food period’ in the annual cycle have been abated; a much larger population of wintering hummingbirds can carry over” (Grinnell and Miller, 1944).Though other introduced plants were also likely involved in the species’ early increase in California, the scale of Eucalyptus planting in the state was truly remarkable. For example, between 1910 and 1914 the Mahogany Eucalyptus and Land Company planted between one and three million Eucalyptus seedlings in the hills lining the east side of San Francisco Bay (O’Brien 2006), creating a near-monoculture forest that blooms abundantly throughout the breeding season of Anna’s Hummingbirds and still characterizes much of the region today.

The first accounts of the species in the northwest were recorded in 1944, both in Oregon (Contreras 1999) and on Vancouver Island (Scarfe and Finlay, 2001). The first northwestern nest report was in 1958 near Victoria, BC (Scarfe and Finlay, 2001). Zimmerman (1973) aggregated reports from birders and breeding bird atlases to document the time of arrivals across the range up to that time, though the species was still considered rare and was not known to regularly breed outside California. Recently, Greig et al. (2017) analyzed a large-scale citizen science dataset of backyard birdfeeder surveys from 1997 to 2013,and documented a range expansion across the northwest seemingly occurring after 1997. They found that Anna’s hummingbirds in the expanded northern range are more likely to visit birdfeeders and occur in human-modified landscapes than those in California, and that changes in climate during the 1990’s and 2000’s are unlikely to explain the observed shifts. Intriguingly, Greig et al. also document a positive feedback cycle between hummingbirds and humans – people who saw hummingbirds were more likely to hang hummingbird feeders, creating an upward pressure on carrying capacity as hummingbird populations increase.

## Methods

### Occurrence Records and Demographic Models

I downloaded records of Anna’s Hummingbird museum specimens from the Global Biodiversity Information Facility (GBIF 2017), and occurrence records from the National Audubon Society’s Christmas Bird Count (CBC; National Audubon Society, 2010) and the Breeding Bird Survey (BBS; Sauer et al. 2017). The combined dataset includes 9,770 occurrence records spanning 1875-2017. To map the timing of arrival in different parts of the range I estimated minimum concave hull polygons for occurrence records in ten-year time bins from 1940-2010, using the “rgeos” (Bivan and Rundel 2013) and “concaveman” (Gombin et al. 2017) packages in R. This analysis was restricted to the first two-thirds of the breeding season (Nov – March) to minimize the signal of post-breeding dispersal. Occurrence records with less than two other records within 100 km were also dropped, on the assumption that they represent vagrant individuals rather than new populations. Concave hull polygons for each period were buffered by 100km to account for dispersal from reported sites. The resulting maps are not meant to capture the absolute first record in each area, but should reflect the time when local abundance rose high enough to be picked up by CBC surveys and museum collectors.

To estimate growth rates and test for density-dependent population regulation, I fit demographic models to CBC data for the period 1950-2016. CBC data was used because the counts occur during the nesting season, the methodology and search areas are standardized, and Anna’s Hummingbirds are strongly associated with the urban and suburban landscapes typically covered in CBC survey circles (Greig et al. 2017). Though many sites report survey results since the 1910’s, data on survey effort is intermittently reported before 1960. Site-years without effort data were dropped from the analysis. As in Soykan et al. 2016, I used party hours as a measure of survey effort, and analyzed the ratio of *C. anna* reports to total survey hours per site/year as an index of abundance.

I removed the highest and lowest abundance index values at each site, subset the data to include only sites with at least 15 years of *C. anna* reports since 1950, and used nonlinear least-squares in R to fit models for each site of (1) exponential growth: *n_i_=n*_0_*e*^*rt*^, (2) logistic growth: 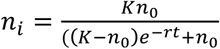 and (3) linear growth: *n*_*i*_=*n*_0_*+ rt*, where *n*_i_ is the abundance in year *i, n*_0_is the starting population size, *r* is the population growth rate, *t* is the number of years since the first record at a site, and *K* is the carrying capacity. Parameter values for the starting population size and carrying capacity were bound by the minimum and maximum observed abundance across all sites. 95% confidence regions for abundance estimates were calculated by generating 1000 bootstrap replicates over survey years at each site, fitting a new model to the bootstrapped data, generating predicted abundances under the bootstrap models, and taking the 95% confidence interval of the predictions. I then ranked models for each locality by Akaike Information Criterion (AIC; Akaike, 1974), taking ΔAIC>2 as moderately strong evidence in favor of the top model. Geographic trends in growth rates and model fits were plotted using the R packages “ggplot2” (Wickham 2016), “maps” (Becker et al. 2017), and “cowplot” (Wilke 2016).

### Climate Niche Modeling

Occurrence records for CBC sites or museum specimens from 1895-1925, 1945-1975, and 1995-2016, were used to train correlative niche models. Each dataset was randomly subsampled to equal size (*n=*452) and cropped to the contiguous United States to match the availability of historic climate data. Interpolated historic climate data from PRISM (PRISM Climate Group 2018) was used to train models on real climate conditions in different time periods. Average monthly precipitation and mean, minimum, and maximum temperatures during the breeding (Nov – May) and nonbreeding (June – October) seasons were used as climate predictor variables. These variables reflect a mix of both climate and life history, because highly vagile animals like birds may make use of different climatic niches during different times of year (Nakazawa et al. 2004). I then used MAXENT (Phillips et al. 2004, 2006) via the R package ‘dismo’ (Hijmans et al. 2017) to fit models to each time period, and examined variable rankings using the built-in permutation test.

To compare niche model output over time, I cross-projected all models – for example, predicting habitat suitability in present climate conditions given a model trained on 1895-1925 climate and occurrence data, and vice versa. I calculated niche overlap *I* (Warren et al. 2008) for all pairs of models and used a Wilcox test to ask if comparisons were significantly different in predictions of the same model across different time periods versus predictions of different models across the same time period. If regional climate change explains most of the range expansion, models trained on 1895-1925 data should predict most of the modern range. Since the range has expanded over time, these models should have low overlap. In contrast if a shift in the species’ realized niche explains the expansion, models trained on a single time period should produce similar projections in all other time periods and thus have high overlap.

Last I tested whether Anna’s Hummingbird climate niches are significantly similar given dispersal limitations and differential availability of climate conditions across time periods using a modified version of Warren et al. 2008’s niche similarity test implemented in R. This test asks whether observed overlap in niche use over time is more or less than what would be expected if a species randomly chose habitat patches within its range. The script first calculates observed niche overlap and estimates available areas for each occurrence dataset by buffering the minimum convex hull of occurrence points by 100km and clipping to coastlines. A null distribution is then generated by randomly distributing points within habitat available in each time period, training models on the randomized dataset, and projecting them to the present. Overlap between observed model projections is then compared to the null of randomized vs observed projections and a *p* value estimated assuming a normal distribution.

### Nesting Phenology

To test for shifts in breeding phenology I assembled records of active nests (females on eggs or later) from natural history museums (via vertnet.org) and eBird (Sullivan et al. 2009). The dataset includes 882 California records, 181 from the southwest, and 124 from the northwest.

I used a Wilcox test to compare median breeding season dates (days since Nov. 1) across native, southwest, and northwest ranges. To ask whether the beginning of the nesting season was delayed, I compared the dates of the 0.1 quantile of breeding days for each region. Last I estimated the impacts of unequal sample sizes by randomly sampling 1000 sets of 124 records from California, recording the difference in medians and tenth quantiles across ranges, then taking the fifth quantile of the resulting distribution as a reasonable lower bound for shifts in timing between regions.

## Results

### Timing of Range Shifts

Museum and CBC data show that Anna’s Hummingbirds were established in the Willamette Valley and Puget Sound by the early 1960’s, and British Columbia by the early 1970’s (Figure 1, Supplementary Figure S1). The first Alaska CBC report is in 1974. These dates are consistent with a 1973 study compiling reports from bird atlases (Zimmerman 1973), but notably later than the first reports by birders on Vancouver island in 1944 (Scarfe and Finlay, 2001). This suggests that the initial phase of colonization occurred during the late 1940’s through the 1950’s, but population densities remained too low to be picked up by standardized surveys. Many early northern records, including the first British Columbia and Alaska reports in this dataset, are from Christmas Bird Counts conducted in December – demonstrating that the range expansion was not restricted to warm months.

**Figure 1.**
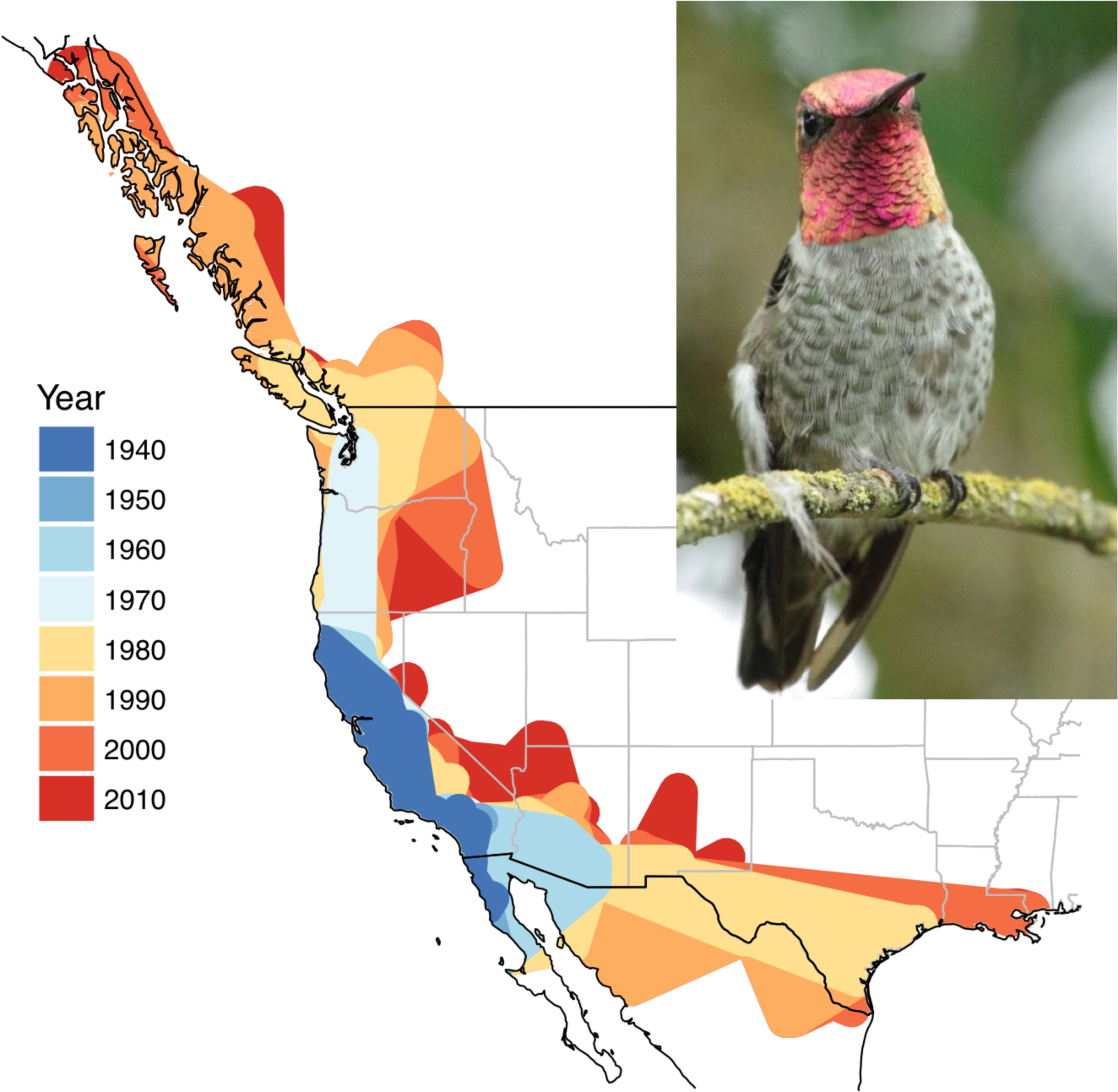
Estimated maximum wintering range (Nov – March) of Anna’s Hummingbirds, 1940-2010. Right: an adult male in Seattle.

Specimens as early as 1925 in Arizona and 1936 in east Texas suggest that the species was either occasional in the region prior to large-scale human development or had already expanded its range by the early 20^th^ century. How-ever, the first wintering records in the southeast are not until 1937 in Arizona and 1967 in Texas, consistent with earlier reports representing post-breeding dispersal rather than local breeding populations. The species quickly expanded across both states during the 1960’s and 70’s, and by the 1980’s was regularly reported across southern Arizona and parts of the Gulf Coast and Rio Grande Valley in Texas.

Several sites in Nevada and New Mexico also reported the species at low frequency starting in the 1970’s. Since 2005 Anna’s Hummingbirds have been regularly reported in Christmas Bird Counts from southwestern Alaska and have appeared as a vagrants as far east as the Atlantic coast of Canada (Supplementary Figure S1).

### Demographic Models

Population growth rates from 1950-2016 range from −0.065 to 0.309 in exponential models (mean .040, 95% quantiles −0.015 to 0.167), with the fastest rates around Puget Sound and the Salish Sea (Figure 2; Supplementary Figure S2-S7). The mean rate estimated here is slightly lower than the estimates of 4.3-5.5% annual growth in Soykan et al. (2016; a comprehensive study of CBC records), suggesting that more sophisticated effort corrections may lead to even higher estimates of population growth.

**Figure 2.**
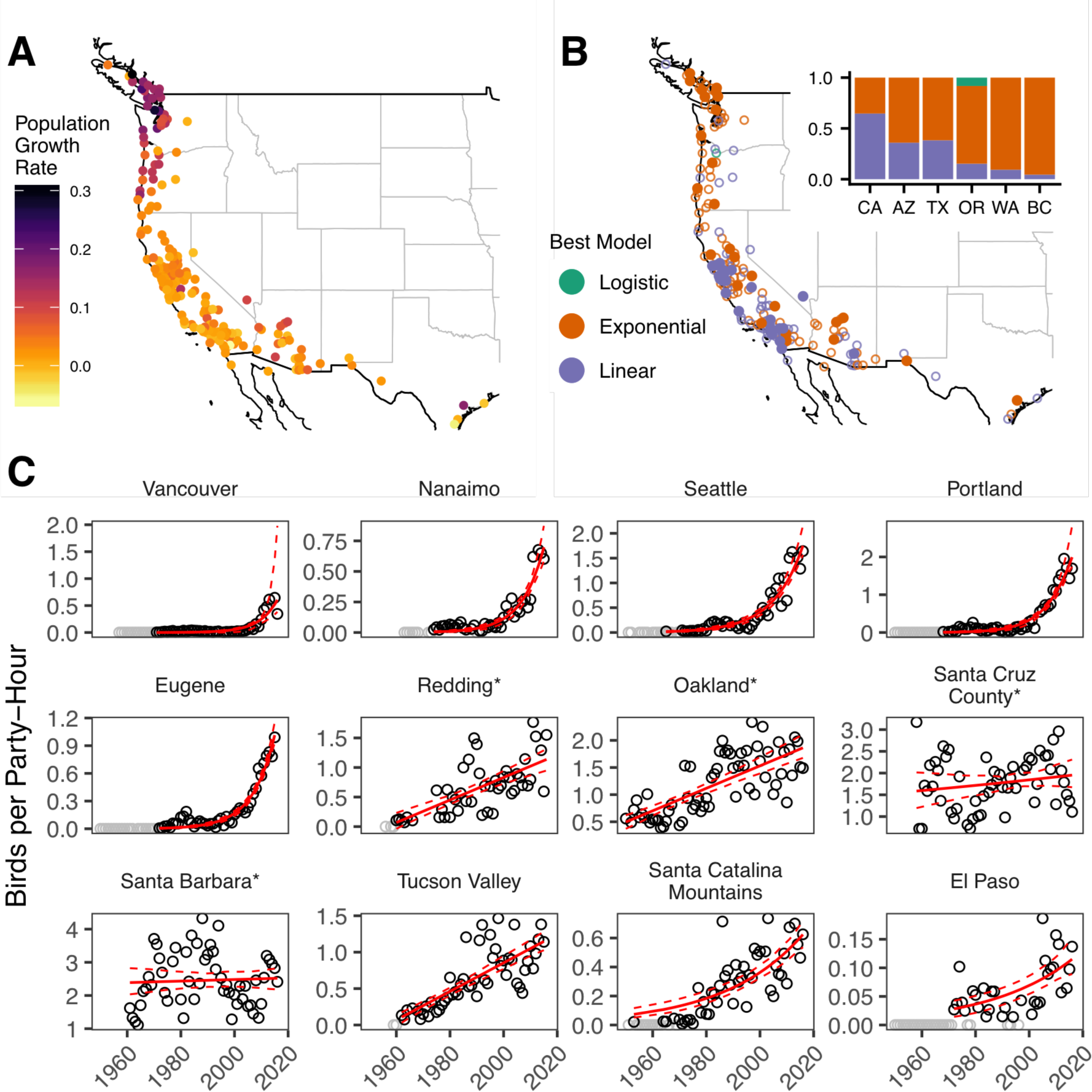
A: growth rate of Anna’s hummingbird populations by CBC count circle. B: AIC model selection by CBC circle. Filled points have a ΔAIC > 2. The inset bars show the proportion of sites with best fits for each model. C: Representative observed (black circles) and predicted (red lines) abundance estimates. Dashed lines are 95% confidence regions and grey circles are years with full survey data but no Anna’s Hummingbird reports. Localities are ordered NW to SE. Asterisks indicate localities in the native range.

In California and much of the southwest, most populations have been growing steadily since CBC effort data became available, with large variability in abundance estimates between years. Sites in Arizona and Texas report Anna’s Hummingbirds starting in the 70’s, with moderate growth rates since colonization. In the northwest populations are consistently reported starting in the mid 1960’s but maintained low densities until the late 1990’s, followed by rapid growth to the present day. There was no relationship between growth rate and colonization time across the full range (linear regression: *p* = 0.729). Growth rates in the northwest were significantly higher than in the native range (Wilcox test: *p*<0.01, 95% CI: 0.08 – 0.11), but those in the southwest were not (*p*=0.176, 95% CI: 0 – 0.02).

Logistic growth models were favored in only 1 of 180 CBC sites across all states. In California, the historic range core, 65% of CBC circles show trends consistent with linear change in population size, while 35% are exponential. In Arizona and Texas the distribution was 55% exponential and 45% linear. In contrast, 88% of sites in the northwest were best described by exponential growth models. Individual model fits in the northwest are surprisingly consistent and show no evidence of density-dependent population regulation. Current densities in northwestern cities like Seattle (1.74 birds/party-hour; bph) and Portland (2.01 bph) have already reached those of cities at the northern half of the historic breeding range such as Oakland (1.92 bph) and San Jose (1.89 bph). Net populations in the north-west; however, are likely still much lower than those in California because the species is more closely affiliated with human-modified habitats in the north (Greig et al. 2017).

### Niche Models

Niche models reflect a shift from a range characterized by nonbreeding precipitation during the early 20^th^ century to one defined by breeding (winter) temperatures after 1995 (Figure 3). This is consistent with Grinnell’s observation that the “minimum food period” late in the dry nonbreeding season historically limited populations (Grinnell, 1944). Niche overlap was significantly higher when comparing projections of the same model across time periods versus projections of models trained in different time periods (Wilcox test, *p<*0.01). Most strikingly, models trained on 1895-1925 data fail to predict any of the high-suitability habitats across the northwest and interior south-west that are currently occupied by expanding breeding populations. This suggests that changes in climate alone cannot explain the range shift. Instead, Anna’s Hummingbirds expanded their realized niche to include colder and wetter habitats.

**Figure 3.**
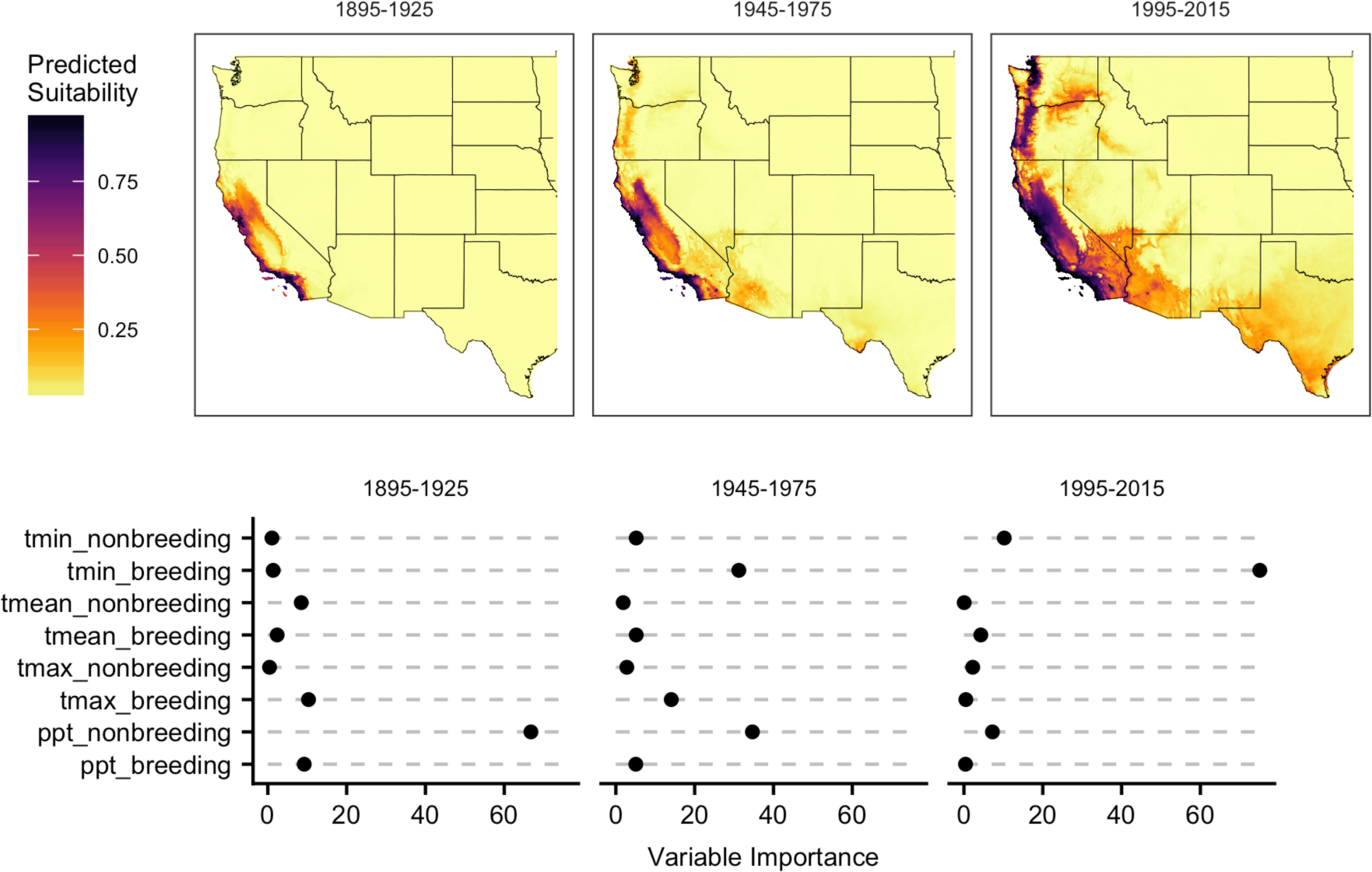
Environmental niche model projections (top) and variable rankings (bottom). All models are projected to 1995-2015 climates.

However, the climate space occupied by Anna’s Hummingbirds was not entirely unrestrained. Niche similarity tests indicated that all models were significantly more similar than expected by chance given random occupation of available habitats (*p <*0.01; observed overlap fell outside the range of the null distribution in all cases). These results show that though the species’ realized niche has expanded, at least some aspects of climate have restrained its distribution across all time periods. High-elevation montane habitats and most of the drier and colder areas east of the Sierra Nevada and Cascade mountains are predicted to be unsuitable habitat in all time periods; likely reflecting innate physiological limits on cold tolerance during the breeding season.

### Nesting Phenology

Nest records show that both the northern and eastern range expansions created new breeding populations (Figure 4). Differences in median nesting date between interior southwest and California records were not significant (*p*=0.146, 95% CI: −2-12 days), but the median nest record in the northwest was 11 days later than in California (*p*=0.009, 95% CI: 7-27 days). The beginning of the nesting season, defined as the tenth quantile of reports, was 25.9 days later in the northwest and 15.9 days later in the southwest. Across randomized equal-sized samples the fifth quantile of differences in medians (a lower bound for the true difference given unequal sample sizes) was −1.5 days for native vs northwest ranges and +2.5 days for native vs. southwest ranges, suggesting that unequal sample sizes could explain observed shifts in the southwest but not the northwest. Lower bounds for differences in the beginning of the nesting season were 18 days for the northwest and 7.7 days for the southwest.

**Figure 4.**
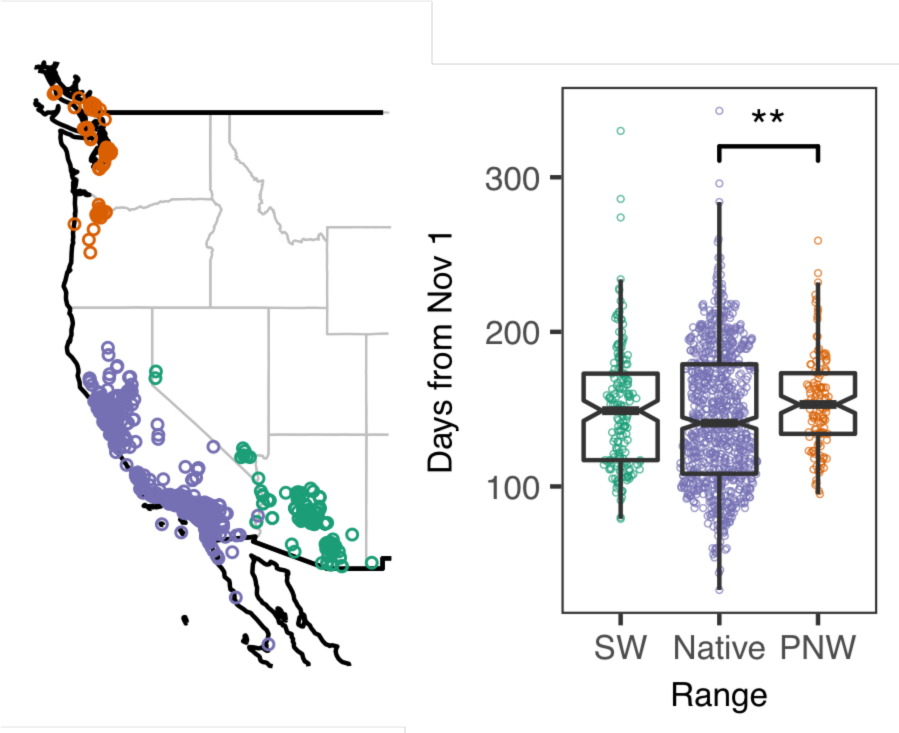
Date and location of active Anna’s Hummingbird nests from museum and eBird data. Asterisks indicate a significant difference.

## Discussion

The Anna’s Hummingbird holds two lessons for studies seeking to predict the magnitude and direction of species range shifts. First, observed climatic associations can be very different from the full niche space in which an organism can live (Peterson, 2003, Waltari et al. 2007, Broennimann et al. 2007). Correlative niche models trained on early records did not predict the species’ modern range, and models trained on recent records found that the current niche is defined by a different mix of variables – in particular, minimum breeding-season temperature – than that in the early 20^th^ century. Grinnell identified Anna’s hummingbirds as close associates of the upper Sonoran life zone, but given the availability of introduced tree and shrub species that flower during the appropriate season the species quickly expanded its realized niche. This niche expansion allowed the species to expand into novel habitats, resulting in exponential growth across the northwest. Despite significant heritability of seasonal timing in at least some birds (Pulido et al. 2001), a mid-winter nesting phenology has also not prevented the species’ ongoing range expansion.

Second, when asking if environmental change explains shifts in range or abundance we should take time lags and demographic process into account (Kowarik 1995, Crooks 2005, Doak & Morris 2010). This means that the dates when we observe what appear to be the largest increases in population size will be later than the environmental or behavioral shifts that allowed the increase in population. It is tempting, for example, to see the sudden uptick in Anna’s Hummingbird populations across the northwest in the 1990’s as evidence that something in the environment changed at that time. However, the consistent fit of northwestern survey records to exponential growth models starting in the 1970’s demonstrates that observed trends can be explained without any change in growth rates or carrying capacity during the 1990’s. Further, niche models constructed from mid-century occurrence and climate data show that the shift towards a climatic niche space defined by minimum breeding temperature was already in progress prior to 1975. If changing behaviors or environmental factors influenced the spread of the species to the northwest, these changes would have occurred in the 1950’s and 1960’s when the first breeding populations were becoming established across the region.

My results support Greig et al.’s 2017 finding that a shift towards more human-modified habitats supported the range expansion in the north, though the longer timespan of survey data here shows that population establishment in the northwest happened during the 1960’s rather than after 1997. The very fast growth observed even in smaller towns and cities in the northwest suggests that supplemental feeding and ornamental plants, rather than the urban heat island effect (Oke 1982), are the most likely direct mechanisms.

Based on historical accounts, niche model results, and the species’ unusual nesting phenology, the most plausible cause of increased abundance up to 1940 is the introduction and extensive planting of winter-blooming trees like *Eucalyptus globulus* in California (Grinnell and Miller 1944, Robertson 1931). It is notable that in addition to those in the novel range nearly all California populations have shown significant growth since the beginning of CBC surveys. The most likely overall model for the range expansion is thus that introduced plants and garden cultivation in California increased population growth rates in the native range, which led to an increase in the number of individuals dispersing across the landscape. This increased population pressure eventually led to colonization of the northwest and interior southwest, where a different set of cultivated plants combined with direct supplemental feeding (i.e. hummingbird feeders) to support new breeding populations in climates that were unsuitable for the species at the beginning of the twentieth century.

How long will populations continue to grow in the northwest? Because current densities in major urban areas like Seattle and Portland have already reached those found in much of northern California, growth will probably level off in these cities over the next decade. If not, we should ask why carrying capacity would be higher outside the native range. Greig et al.’s finding of an increase in dependency on hummingbird feeders in the north could provide a likely answer. This theory could be tested by comparing isotope ratios from hummingbirds in the northwest and native ranges – particularly because many hummingbird feeders are filled with a sucrose solution made from sugarcane, a grass which should contain the characteristic signal of C4 photosynthesis in its carbon atoms (Schirnding et al. 1982). The species is also a promising subject for genetic studies of the impacts of the impacts of range shifts, both because the demography is relatively well known and because a high quality genome already exists due to its use as a model system for the song learning (Korlach et al. 2017).

The Anna’s Hummingbird has become a common urban breeding bird across the Pacific Northwest by expanding its realized climate niche, shifting its unusual nesting phenology to account for colder and wetter winters, and likely by increasing its association with human-modified habitats (Greig et al. 2017). This range expansion appears to be the product of increased local abundance within the native range driven by population growth and plant introductions in the mid nineteenth century. Although direct supplemental feeding is uncommon for most species, nearly all ecosystems are subject to human modification of the environment via the introduction of new plant species, and historical records coupled with niche model results suggest that this factor was important both in increasing early populations in California and allowing the species’ colonization of the northwest. Whether the invasion dynamics observed in Anna’s Hummingbirds are typical of range shifts of native species during the twentieth century remains to be seen, because many recent studies have been limited to binary presence/absence grids (Tingley et al. 2009, Parmesan et al. 1999, Chen et al. 2011, Greig et al. 2017). As the magnitude of available survey data increases over time our ability to infer changes in both presence and abundance should increase across many taxa, allowing a more mechanistic view of how, rather than whether, species’ ranges change over time.

## Data Availability

A summary of model parameters by site for the CBC dataset and R scripts used for analyses and figures are available at: https://github.com/cjbattey/anhu. Full CBC data can be requested from.

## Acknowledgements

Many thanks to Audubon Society staff and the roughly 270,089 volunteer field counters who participated in the Christmas Bird Counts analyzed here. Ethan Linck, Ray Huey, Cooper French, Dave Slager, and John Klicka provided helpful comments on earlier versions of this manuscript. This project was partially supported by NSF Grant #DEB-1600945.

“Because of human settlement of open valleys and plains and the clearing of woodland, with extensive gardening and the planting of flowering, non-native trees, the numbers of Anna Hummingbirds now no doubt greatly exceed those comprised in the original aggregate population.”

—Grinnell & Miller, 1944

